## Supplementary material for "Tissue libraries enable rapid determination of conditions that preserve antibody labeling in cleared mouse and human tissue": Public peer review and response to reviewers

Dear Editors of eLife,

We would like to thank you for the very constructive and positive critique from the first round of peer review. We have made significant efforts to address all of the suggestions for improvement including two new sets of experiments and 8 new supplemental figures of quantitative information requested by reviewers, as well as a multitude of changes including 1) additional experiments that quantify the saturation of antibody saturation at different time points and incubation temperatures, 2) additional experiments that label multiple brain regions to test the assumption that antigens have a relatively constant distribution with the depth of imaging, as well as methodological improvements for situations when this assumption is incorrect, 3) re-analyzed new and old data using the machine-learning software, Ilastik, and include quantification of signal intensity distribution, background signal distribution, and additional performance parameters such as the distance into the tissue at which the staining intensity drops by half. Analyses with these new methods show the same trends in staining quality for different conditions as reported in our initial manuscript, but the segmentation provided by ilastik is more robust than before which has the benefit of reducing variation between replicates. We were also pleasantly surprised that ilastik was free to use and enabled batch segmentation, so we wrote macros for ImageJ to convert data to the proper file formats that facilitate the batch analysis of the entire library at once in ilastik and can feed into Matlab to quickly provide the signal, noise, and total segmented area as a function of depth through the sample as well as the distance into the tissue at which the staining intensity drops by half. This significantly cuts the time and labor involved in the analysis of these data, and its free cost will allow for others to more easily adopt our methodology for immunohistochemistry optimization, so we are extremely grateful to reviewer 1 for suggesting this software.

In addition, we've responded to each comment and made changes in the manuscript as described below:

### Reviewer #1:

#### Tissue thickness and staining time choices should be justified more carefully. Of particular concern is the antibody staining time, as the optimal time depends strongly on the temperature of incubation. Staining should be quantified at several time points to determine the extent of saturation at the tissue surface.

We have conducted additional experiments to compare room temperature and 4$^{o}$C incubation of NeuN at 5 time points, plotted in figure **S1**. These results show both the increase in saturation at the tissue surface and within the tissue. For both incubation temperatures, the tissue appears to saturate at the surface within 24 hours but there are significant differences in the staining intensity with depth. Neither shows significant signal at a depth of 250 microns, suggesting that 500 micron thick tissue is adequate for comparing these conditions with <24 hour incubation times.

#### The assumption that antigens have a "relatively constant distribution with the depth of imaging" should also be supported with data. The same goes for lateral variation. Variation between replicates also would ideally be reduced.

We conducted additional experiments to compare NeuN and Glut1 staining across three different brain regions in longitudinal slices from each hemisphere of three different mice. We incubated these antibodies for 3 days to achieve saturation, then quantified the average signal, percent segmented area, average noise, and signal/noise ratio in each imaging plane. Our original metric of percent segmented area shows some variation with depth depending on the brain region, but we find that normalizing the signal intensity per segmented area allows for a quantifiable metric for staining quality that is independent on the change in distribution of antigen, so long as there is some antigen in each plane. These data show insignificantly different results across tissue samples and brain regions. Additionally, as we will describe below, Ilastik segmentation is more robust which decreased a significant amount of variation between replicates. We feel the variation between replicates is reasonable, as quantified by the Signal-Noise ratio standard error of mean values in the hippocampus (10.2±2.4%) anterior cortex (11.0±3.5%) and posterior cortex (10.3±1.8%), especially when considering these tissue samples came from different mice.

**From the methods, it seems as though the antibody is simply added to the staining buffer with the tissue, which might indicate there is insufficient mixing before and during antibody**

We have clarified in the methods section that the antibody was mixed in the staining buffer, and that the tissue was placed on a gentle shaker during antibody incubation for all conditions.

**Image analysis: A more sophisticated but still highly user-friendly segmentation strategy such as the Ilastik software package might be a more reliable and versatile choice for analyzing image data.**

We attempted to use Ilastik, as recommended, and found it to be an extremely robust solution for analyzing our data. Before, using Imaris, we did not have access to the batch processing module due to its cost. This meant that we had to go through each file and individually generate the segmentation mask for each channel, which was a significant amount of time for the hundreds of imaging files used in this work. Ilastik allowed for us to train the segmentation on a subset of our data, then apply the segmentation to all of the similar images. Additionally, this did not require the normalization step we had applied to our data in Imaris, further improved the segmentation of poorly stained samples, and allowed for the isolation of nonspecific staining that may have confounded analysis. We are extremely grateful for this suggestion.

**Furthermore, the signal intensity distribution within the masked signal area can be extracted (e.g. in Matlab) to quantitatively compare processing conditions, in addition to the area occupied by the mask as used here. The area outside the mask could potentially be treated as a background segment, whose intensity distribution can also be quantified. This way, quantitative measures of both signal intensity and background staining levels could be used for a more full comparison between method**

**variants. The results should be viewed as a 1D line plot for more quantitative comparison between conditions. Finally, this lineplot could be used to measure derived performance parameters such as the distance into the tissue at which the staining intensity drops by half.**

We appreciate these very constructive and insightful suggestions. As suggested, we performed this analysis on four of our sets of data (Mouse NeuN, Mouse Glut1, Human NeuN, and Human Glut1) consisting of 142 image volumes as a proof-of-principle. These quantifications provide some additional insight into the difference between conditions. Most interesting is the significant difference in the percent staining area in each imaging plane was more changed than the signal intensity across conditions, which provides support for our claim that IHC is significantly influenced by denaturation or masking of the epitope. The antibody is having the same overall effectiveness for signal intensity per unit area, but is binding to less. Additionally, the performance parameter suggested by the reviewer (which we call half-max depth in the manuscript) showed reproducible differences across samples and translated well between mouse and human tissue. We included this plot in Figure 1 and Figure 5.

### Reviewer #2:

**Results from a small number of antibodies are extrapolated to apply to all antibodies in general.**

Although we have used relatively few antibodies compared to the many thousands used in research, we rigorously tested each of the antibodies in this study. We have collected a large amount of data for each antibody, which includes at least three replicates from independent brains for 12 different conditions. Additionally, each replicate includes approximately 100 imaging planes with 2-4 channels, depending on the stains that were tested. We feel confident that these data strongly support our main claims that small changes in conditions lead to significant variation in IHC quality, which we see among the antibodies we tested. Additionally, the variation in IHC quality for a dozen different conditions is consistent between mice and human tissue that are prepared similarly. These results are what allowed for us to find conditions that produce the beautiful images in figure 6 that show simultaneous staining of multiple antibodies in human tissue. That said, we did not rigorously test the use of conjugated primary antibodies compared to use with secondary antibodies with multiple antibodies, so we have removed those claims which we feel are a minor point in the original manuscript.

**The authors conclude that "IHC is limited more by denaturation of the epitope than the antibody itself". This conclusion is based on the experiments that show that varying the conditions of tissue preparation and staining can dramatically change antibody performance from no label to excellent label. However, the authors do not take into account that they are using antibodies that have already been shown to work reasonably well for immunolabeling.**

We believe that the new analyses suggested by reviewer 1 help support our claim that IHC may be limited more by the denaturation of the epitope than the antibody itself. We agree that we are using antibodies that have already been shown to work reasonably well for immunolabeling, and have included mention of this in the second paragraph of the results section.

**Important methodological information is missing in some instances, for example how was the human tissue fixed or how are the tissue libraries stored.**

We have included more details of our methodology in the *Human tissue samples* and *Immunohistochemistry* sections, as well as added a section on the *Storage of tissue library*

**Results, Measuring IHC quality in cleared tissue: It is unclear how the threshold was determined. Another concern is that this "measure of IHC quality" is essentially a measure of what fraction of the tissue is labeled. Thus, it does not account for specific vs non-specific staining.**

We believe that the new methodology using ilastik addresses reviewer 2's concerns as well. Using this new methodology there is no need for setting a threshold. It does require the users to properly train the segmentation software, however the results are updated live during training which allows for users to quickly assess its quality. Furthermore, you can train ilastik to separate non-specific staining from the signal and background which allows for quantification of non-specific staining.

**Conditions that influence antibody and epitope preservation: It is unclear whether the baseline protocol described here applies only to mouse tissue or to human tissue as well. The mention of perfusion leads me to believe this is specifically for mouse, but I couldn't find information on how the human tissue was fixed.**

We have included more details of this methodology in the *Human tissue samples* and *Immunohistochemistry* sections.

**There is no mention here of whether the antibodies (GluT1 and NeuN) are directly conjugated to fluorophores. I assumed that they were, based on the info in the Methods section. But then in the last paragraph of Results the authors say that "Therefore, we directly conjugated three antibodies to three different fluorophores (NeuN-A568, Glut1- A488, and AT8-A647)."**

We have included more details of this methodology in *Immunohistochemistry* section, as well as in the figure 1 and 5 captions.

**Comparison of IHC conditions for different antibodies directed against a single protein: As explained in the Public review section, I disagree with the conclusion "it is recommended to use primary antibodies directly conjugated to fluorophores whenever possible." Unless the authors add more experiments that compare direct vs. indirect immmunolabeling with the same primary antibody (for several different antibodies), this conclusion is not justified.**

As recommended by reviewer 2, we have withdrawn the conclusion that our data shows significant advantages of primary over secondary antibodies for reducing nonspecific staining.  This would take a significant amount of effort for a relatively minor point, which we may be able to address in the future, however we are currently exclusively using conjugated antibodies in all our experiments. We have still included a recommendation to use conjugated antibodies where possible because it reduces the number of steps and total time for IHC, sometimes on the order of weeks depending on tissue thickness.

**"In order to examine how much of the observed variability in antibody labeling is due to targets being more or less accessible depending on cell type or subcellular location, multiple antibodies were also chosen that recognize tau protein." I don't understand what the comparison of the tau antibodies will tell us about the accessibility of targets depending on cell type or subcellular location.**

We have removed our statement on subcellular location, but hope that we can address this in future work.

**Mouse brain tissue as a model system for optimizing IHC in cleared human tissue: Determining the optimal protocol for simultaneous staining of 3 antibodies is an important point. It will be very useful if the authors elaborate more on how this optimal protocol was determined. How was the combination of conditions that maximize staining quality established?**

We have clarified our process for determining the optimal protocol from the collected tissue library data in the main text in multiple places, including the following text under **Mouse Brain tissue as a model system for optimizing IHC in cleared human tissue** in the results section: "The optimal protocol that allows NeuN, Glut1, and AT8-tau staining was then determined by finding the conditions that allow for the simultaneous IHC of all three antibodies in a single tissue section, and then finding the combination of conditions among those that maximizing the staining quality. It is worth noting that this was not necessarily the best protocol for each individual antibody. It is more important to have adequate staining of each epitope than to have fantastic staining of one at the cost of another. As described above, we determined the optimal protocol for multiplexed labeling by combining the protocol changes that quantitatively improve the staining quality, defined by half-max depth, for each antibody above the baseline protocol and do not significantly degrade performance of other antibodies. For NeuN, Glut1, and AT8 antibodies this protocol involves clearing for 2 days, using 50$^{o}$C to unmask epitopes, and incubation of antibodies at room temperature."

**Minor point: how was the 14 days duration determined?**

Admittedly, the 14 days duration was due to the antibody incubation occurring over a holiday and generally we have found that there is no problem with longer incubation times above what is necessary. In newly collected data incorporated with the time-series (Figure S1), we have shown the minimum time for saturation of a similar tissue slice is on the order of 3 days.

**Discussion: "Rather than the antibody, our data suggests that poor staining is most often due to denaturation or blockage of the epitope." I disagree with this conclusion, as explained above.**

We believe that the new analyses suggested by reviewer 1 help support our claim that IHC may be limited more by the denaturation of the epitope than the antibody itself. We agree that we are using antibodies that have already been shown to work reasonably well for immunolabeling, and have included mention of this in the second paragraph of the results section.

**"Unfortunately, as we found with the dopamine receptor 2 and GFAP IHC, a protocol may not exist to allow for both targets to be effectively stained simultaneously." - I didn't see this presented in the Results section.**

It is described in the section **Conditions that influence antibody and epitope preservation** in the second paragraph, specifically: "Inclusion of PFA in the hydrogel resulted in decreased IHC quality for both Glut1 and NeuN, however this significantly improved IHC of dopamine receptor 2 (figure S11)."

**How were the human samples fixed? Were the slicing and all the following steps performed in an identical way? I assume they were, but this should be stated clearly in the methods. How were the tissue libraries stored? For how long can they be used after preparation?**

We have added a new section "Storage of tissue library" with the following text: The tissue library was then used immediately or stored at 4°C in PBS with 0.02% (w/v) sodium azide. Mouse and human tissue was processed following all of the steps above and then used up to 3 months later. Additional mouse and human tissue was stored following delipidation, then successfully used up to 6 months later by resuming from the pre-immunohistochemistry tissue processing state. IHC conditions for a given antibody were always compared using batches of tissue that were stored for the same amount of time.

**Figures 1, 2 and others. How is the top of the side view determined? Is this the surface of the tissue and therefore conditions where the label in the side view starts lower that in other conditions means that the tissue surface was not labeled? And why would that be? If that is not the case, it will be best to align the side views and have the labeling start at the same level.**

We have aligned the side views as recommended by reviewer 2. In these images, sometimes the imagine plane started slightly above the top plane of the tissue. For the purposes of quantification and presentation, our matlab code now starts the plot from the first plane with 80% of the max staining intensity within a sample. This is clarified in the materials and methods where we describe the code for analysis. The top view is chosen arbitrarily as a section that faithfully represents the quality of the rest of the image.

**Figure S3. This is a very clear way of summarizing the differences in labeling. It would be very helpful to have this summary next to the actual immunostaining images in the main figures.**

We have added the half-max intensity values to figure 1 near the actual immunostaining images to summarize the differences in labeling.

### Figure S9 Representative microtome images taken from the human tissue prior to staining: Do you mean prior to clearing?

In figure S9 (now figure S15) we do mean that the images were taken prior to clearing, and have changed the text to clarify that point.
